# Engineering A Fluorescent Protein Color Switch Using Entropy-driven Beta Strand Exchange

**DOI:** 10.1101/2021.10.20.465183

**Authors:** Anna M. John, Harsimranjit Sekhon, Jeung-Hoi Ha, Stewart N. Loh

## Abstract

Protein conformational switches are widely used in biosensing. They are typically composed of an input domain (which binds a target ligand) fused to an output domain (which generates an optical readout). A central challenge in designing such switches is to develop mechanisms for coupling the input and output signals via conformational change. Here, we create a biosensor in which binding-induced folding of the input domain drives a conformational shift in the output domain that results in a 6-fold green-to-yellow ratiometric fluorescence change *in vitro*, and a 35-fold intensiometric fluorescence increase in cultured cells. The input domain consists of circularly permuted FK506 binding protein (cpFKBP) that folds upon binding its target ligand (FK506 or rapamycin). cpFKBP folding induces the output domain, an engineered GFP variant, to replace one of its β-strands (containing T203 and specifying green fluorescence) with a duplicate β-strand (containing Y203 and specifying yellow fluorescence) in an intramolecular exchange reaction. This mechanism employs the loop-closure entropy principle, embodied by folding of the partially disordered cpFKBP domain, to couple ligand binding to the GFP color shift. This proof-of-concept design has the advantages of full genetic encodability, ratiometric or intensiometric response, and potential for modularity. The latter attribute is enabled by circular permutation of the input domain.

## Introduction

Protein switches are versatile tools with applications in synthetic biology, molecular diagnostics and cellular biosensing ^1–4^. Rapid advancements in the field of protein switch engineering using random, rational, semi-rational, and *de novo* approaches testify to the ongoing search for improved designs^5–8^. Many of these efforts aim to achieve two key goals: *signal transduction* (i.e., a strategy for joining recognition and reporter domains such that the signaling event is transduced to a detectable output) and *modularity* (the capability of substituting different recognition domains for sensing a target of choice)^9,10^. In this study, we address these challenges by employing a combination of two protein engineering strategies — loop closure entropy^11^ and alternate frame folding (AFF^12^) — to create a conformationally-driven biosensor in which ligand binding and change in fluorescence color/intensity comprise the respective input and output signals.

AFF is a mechanism by which allostery is introduced into a protein which previously had none. AFF entails copying a terminal segment of a protein that contains a key functional residue, e.g., one that binds a target ligand, is essential to enzymatic function, or specifies light absorption/emission wavelengths, and pasting it to the opposite terminus of the parent protein. The presence of two identical segments allows the protein to switch between two folding ‘frames’, one of which corresponds to the original amino acid sequence and the other to that of a circular permutant (CP). The two folds are mutually exclusive because the duplicated segments compete for the shared region. Changing the identity of the key residue in one of the segments links the fold shift to a change in protein function. We previously applied the AFF methodology to proteins that bind calcium (calbindin D_9k_ ^13,14^) and a sugar (ribose binding protein_15_), to convert them to biosensors for their respective ligands. Förster resonance energy transfer (FRET) output was achieved by attaching chemical fluorophores to sites that reported on the fold shift.

Here, we introduce two new features to the AFF design: full genetic encodability and a novel mechanism (loop closure entropy) for transducing ligand binding to a fluorescence change. We demonstrate proof-of-concept by creating a genetically encodable protein switch that functions as a small molecule-induced ratiometric biosensor *in vitro* and an intensiometric biosensor in mammalian cells. This biosensor is designed to be generalizable by inserting different recognition domains into the same fluorescent protein (FP) scaffold.

## Results

### Biosensor design

The biosensor color change is made possible by switching between two amino acids at position 203 in strand β10 of the 11-stranded β-barrel of a modified Clover GFP^16^ (referred to as GFP in this paper; amino acid sequences of all constructs are shown in Figure S10). Residue 203 contacts the chromophore (G65-Y66-G67) that resides between β3 and β4. Replacing T203 with Y203 shifts the emission maximum from 509 nm (green) to 524 nm (yellow)^17,18^. To enable the AFF conformational change, we employed the design introduced by Boxer and colleagues to construct a light-driven, fluorescent protease sensor^19^. GFP was first engineered to terminate its sequence with β10 instead of β11. This was achieved by circularly permuting it between β10 and β11 and joining the original N- and C-termini with a GGGSGG linker (Figure 1A). We then appended a duplicated β10, carrying the Y203T mutation, to the N-terminus of the CP to generate the FP scaffold (Figure 1B). The resulting fusion protein can adopt either the green fold (G-fold; GFP permuted between β10-β11 and with T203) or the yellow fold (Y-fold; GFP permuted between β9-β10 and with Y203). The conformational change consists of the duplicate β10 strands exchanging from the common core of (β11, β1-β9) (Figure 1A).

**Figure 1.**
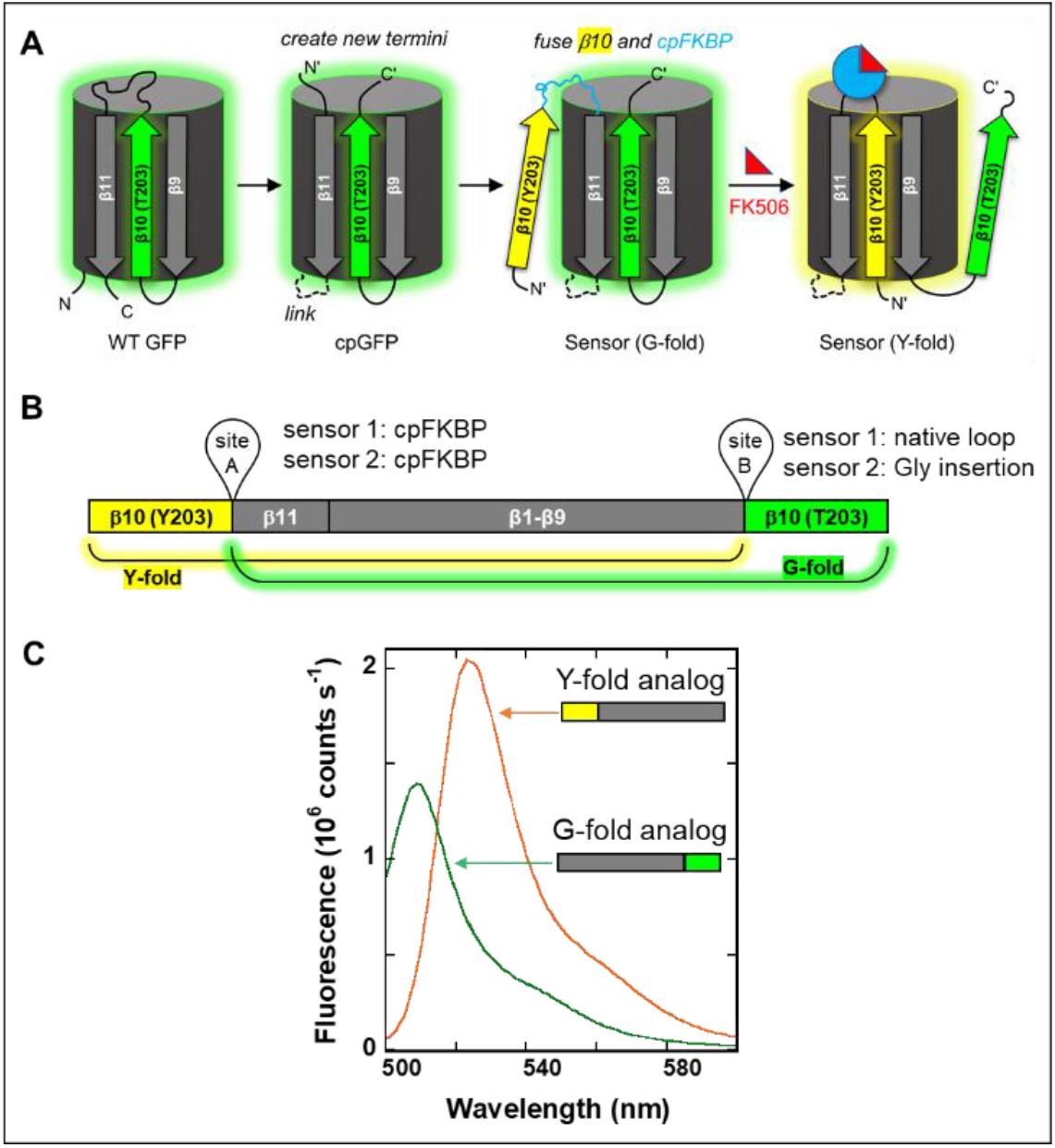
Design of esAFF sensors. (A) Sensors 1 and 2 were created by circularly permuting GFP at the loop between β10 (green, with T203) and β11, and joining the original termini with a 6-amino acid linker (dashed loop). The new termini are designated N’ and C’. A duplicate β10 strand (yellow, with Y203) and cpFKBP (blue) were then appended to N’, establishing the mutually exclusive G-fold and Y-fold. The grey β-strands and cylinder represent the structure shared between the G-fold and Y-fold. In the absence of ligand, the cpFKBP domain is mostly disordered and the sensor populates the G-fold. Upon binding FK506 or rapamycin, the cpFKBP domain becomes structured and stabilizes the Y-fold. (B) The linear depiction of the esAFF sequence shows the locations in which the receptor domain (site A) and thermodynamic tuning peptides (site B) were inserted to generate sensor 1 and sensor 2. Color schemes are identical to those in panel A. (C) Fluorescence spectra of the isolated G-fold and Y-fold analogs recorded at the same protein concentration shows that the Y-fold is 1.675 times brighter than G-fold as defined by the integrated areas-under-the-curve (excitation at 492 nm). Insets show the G-fold and Y-fold analogs indicating the positions of β10 with T203 (green) and Y203 (yellow).

We linked the fold shift to molecular recognition using the principles of loop-closure entropy and binding-induced folding. Loop-closure entropy refers to the loss of chain entropy that results from constraining the termini of a disordered, flexible polypeptide to a single, fixed position^11,20^. When a disordered recognition domain is inserted into a host protein, the host pays the entropic cost of constraining the termini, and the host becomes destabilized. This penalty goes away when the ends become fixed by another process, e.g., binding-induced folding. If the distance between the termini of the folded recognition domain is compatible with the structure of the host protein at the insertion point, the host protein becomes stabilized by ligand binding.

To drive color change of the FP scaffold in Figure 1B, one can insert the disordered recognition domain (circularly permuted FK506 binding protein, or cpFKBP) into one of two positions that will selectively destabilize the Y-fold and not the G-fold or vice versa. These sites are the turn between β10-β11 in the Y-fold (site A in Figure 1B) or the turn between β9-β10 in the G-fold (site B in Figure 1B), respectively. On binding FK506 or rapamycin, folding of the cpFKBP domain removes the entropic penalty and stabilizes the fold in which it is resident. We denote this switching mechanism as entropy switching AFF (esAFF).

### Characterization of sensor components

We first purified and characterized the individual proteins from which the FP scaffold was assembled: the isolated G-fold analog and the isolated Y-fold analog (Figure 1C, inset). Spectral characterization confirmed the expected absorbance maxima (492 nm and 514 nm; Figure S1A) and emission maxima (509 nm and 524 nm; Figure 1C) of the G-fold analog and Y-fold analog, respectively. As anticipated from previous studies^18^, Y203 increased the brightness of the chromophore: emission of the Y-fold analog was 1.675-fold greater than that of the G-fold analog when both proteins were excited at 492 nm, as determined by integrating the areas under the curves (AUC) from 500 to 600 nm for each analog’s emission spectra (Figure 1C). Binding of FK506 to the Y-fold analog had no effect on its fluorescence (Figure S1B). Guanidinium hydrochloride (GdmCl) denaturation experiments revealed that the Y-fold analog is more stable than the G-fold analog as judged by the apparent change in the Gibbs free energy of folding (ΔΔG_fold_ = 2.78 kcal/mol) and by the change in midpoint of denaturation (ΔC_m_ = 0.80 M GdmCl) (Figure S1C; Table S1).

The criteria for the target recognition domain in the esAFF design are that it: (i) has a short N-to-C terminal distance (less than or equal to the C_α_-C_α_ distance between the residues at the ends of the loop into which it’s inserted); (ii) is predominantly unfolded in the absence of ligand; and (iii) folds on ligand binding. To shorten the 26 Å N-to-C length of WT FKBP, we circularly permuted it at position 33. Conveniently, cpFKBP is ∼50 % unfolded without ligand, and fully folds upon binding FK506 or rapamycin (Figure S2A). Binding and folding occur within the dead time of mixing (∼10 s); Figure S2B).

To create the esAFF biosensor, cpFKBP can either be inserted into the Y-fold or the G-fold (site A or site B; Figure 1B). We chose the Y-fold because it is more stable than the G-fold analog. As expected, insertion of cpFKBP destabilized the Y-fold analog, as evidenced by the substantial decrease in C_m_ (ΔC_m_ = 1.83 M GdmCl) (Figure S1C; Table S1). This finding lends support to the loop closure entropy model. We could not, however, quantify this effect because the cooperativity parameter (m-value) significantly increased (∼2-fold) after cpFKBP insertion (Table S1), complicating the interpretation of ΔΔG_fold_. Likewise, we were unable to measure the extent to which FK506 binding/cpFKBP folding stabilized the Y-fold analog because the cpFKBP/FK506 complex unfolds at lower [GdmCl] compared to the Y-fold analog.

### In vitro sensor performance

To quantify color changes, we unmixed the sensor emission spectra into linear combinations of green and yellow reference spectra (obtained from Figure 1C; see Methods). Experimental spectra were reproduced adequately by the sum of the unmixed green and yellow areas-under-the-curve (AUC_green_ and AUC_yellow_) with no other fluorescent species detected (representative examples are shown in Figure S3). The yellow-to-green fluorescence ratio (Y/G) was defined as AUC_yellow_/AUC_green_, with (Y/G)_ON_ and (Y/G)_OFF_ corresponding to the values with ligand or DMSO vehicle, respectively. The turn-on ratio—the main measure of sensor performance—was then calculated by dividing (Y/G)_ON_ by (Y/G)_OFF_.

We also considered the possibility that the fluorescence change could be due in part to an AFF-independent mechanism, e.g., ligand binding changing the protonation state of the mature chromophore or causing the chromophore to mature. Whether such processes contribute to the observed turn-on can be determined by comparing the increase in concentration of Y-fold to the decrease in concentration of G-fold, by adjusting the ratio of their AUC values with their known difference in brightness (see Methods). In other words, if AUC_yellow_ increased by more than 1.675 times the value by which AUC_green_ decreased, then the excess amount was attributed to additional chromophore formation in the Y-fold.

Sensor 1 was constructed by inserting cpFKBP into site A of the FP scaffold as shown in Figure 1B. We expected sensor 1 to express mostly in the G-fold because the G-fold analog was more stable than the Y-fold analog (Table S1). Surprisingly, the purified protein exhibited a (Y/G) value of 2.65, indicating that it expressed mostly in the Y-fold (equal populations of Y-fold and G-fold would result in (Y/G) = 1.675). We hypothesized that the Y-fold might be favored due to a kinetic folding trap, as proposed by Boxer *et al*.^19^. In their design as well as ours, the Y203-containing β10 strand is the first to emerge from the ribosome and the T203-containing β10 strand is the last to exit. Since co-translational folding of GFP is efficient^21^ and chromophore residues interact more favorably with Y203 than they do with T203^19^, it may potentially introduce a kinetic trap for the Y-fold. As a test, we unfolded sensor 1 with HCl (pH 2) and refolded by rapid pH jump back to pH Sensor 1 refolded mostly in the G-fold ((Y/G) = 0.13, Figure S3A), consistent with the kinetic trap hypothesis. We applied this unfolding/refolding procedure before testing all sensors *in vitro*.

Addition of FK506 to sensor 1 failed to produce a significant turn-on (Figure S3B). The likely reason is that the thermodynamic balance between the G-fold and the Y-fold is too far in favor of the former to be overcome by loop-closure entropy gains to the latter. To bring them into closer balance, we sought to selectively destabilize the G-fold using the same loop entropy strategy. We created G-fold analogs with 1-mer (Gly), 5-mer (GSGSG), and 10-mer (GGSGTSGGSG) unstructured peptides inserted into site B (Figure 1B). The single Gly insertion was sufficiently destabilizing, making the apparent stability of the resulting G-fold analog (G-fold analog 2) approximately equal to that of the Y-fold (Figure S4). Increasing insertion length to 5 and 10 residues produced only minor additional decreases in stability (Figure S4). We therefore constructed sensor 2 by inserting a single Gly into site B (Figure 1B). As expected, destabilizing the G-fold increased the (Y/G) value of the purified protein to 24, and that of the refolded protein to 0.86 ± 0.02 (Figure S5), the latter value being consistent with the near-equivalent stabilities of the G- and Y-folds.

We evaluated the performance of sensor 2 at 22 °C, 37 °C, and 45 °C. Visual inspection of sensor 2 emission spectra confirmed a robust shift from green to yellow fluorescence upon FK506 addition at 45 °C (Figure 2A). The time-dependent spectral change resembled that of the theoretical shift from OFF to ON states, as visualized by the superposition of the isolated G-fold and Y-fold spectra (Figure 1C). Unmixing the sensor 2 emission spectra revealed a (5.97 ± 0.83)-fold turn-on after FK506 exposure, relative to the DMSO vehicle control (Table 1). To measure the rate of the conformational shift in the presence of ligand, we fit the time-dependent changes in AUC_green_ and AUC_yellow_ to single exponential functions (Figure 3B). Both half-times (t_1/2_) were approximately equal at ∼ 5 h (Table 1). The switching rates appeared to slow down with temperature, to the point where little change was observed at 22 °C or 37 °C over the course of 20 h (Table S2). To determine reversibility, we incubated sensor 2 with 15 µM FK506 for 9 h (Figure S7A), and then added 30 µM WT FKBP. No significant change in Y/G ratios were observed after 40 h, indicating that the conformational change was irreversible on this time scale (Figure S7B).

**Table 1.** *In vitro* characterization of esAFF sensors^a^.

| Sensor ([GdmCl]) | Turn-on | $t_{1/2}$ (h) Y-fold | $t_{1/2}$ (h) G-fold | % Turn-on due to AFF |
| --- | --- | --- | --- | --- |
| 1 | 1.11 | NA | NA | NA |
| 2 | $5.97 \pm 0.83$ | $4.16 \pm 1.1$ | $5.76 \pm 0.43$ | $89.8 \pm 15.5$ |
| 2 (0.15 M) | $6.64 \pm 1.41$ | $5.58 \pm 2.06$ | $5.52 \pm 1.6$ | $99.3 \pm 8.9$ |
| 2 (0.3 M) | $6.32 \pm 0.59$ | $6.39 \pm 1.40$ | $5.73 \pm 2.72$ | $110 \pm 5.5$ |

<sup>a</sup>The values represent the average ± SD ( $n$ = duplicate or triplicate). In the table, esAFF and NA denote entropy-switching alternate frame folding and not applicable, respectively.

**Figure 2.**
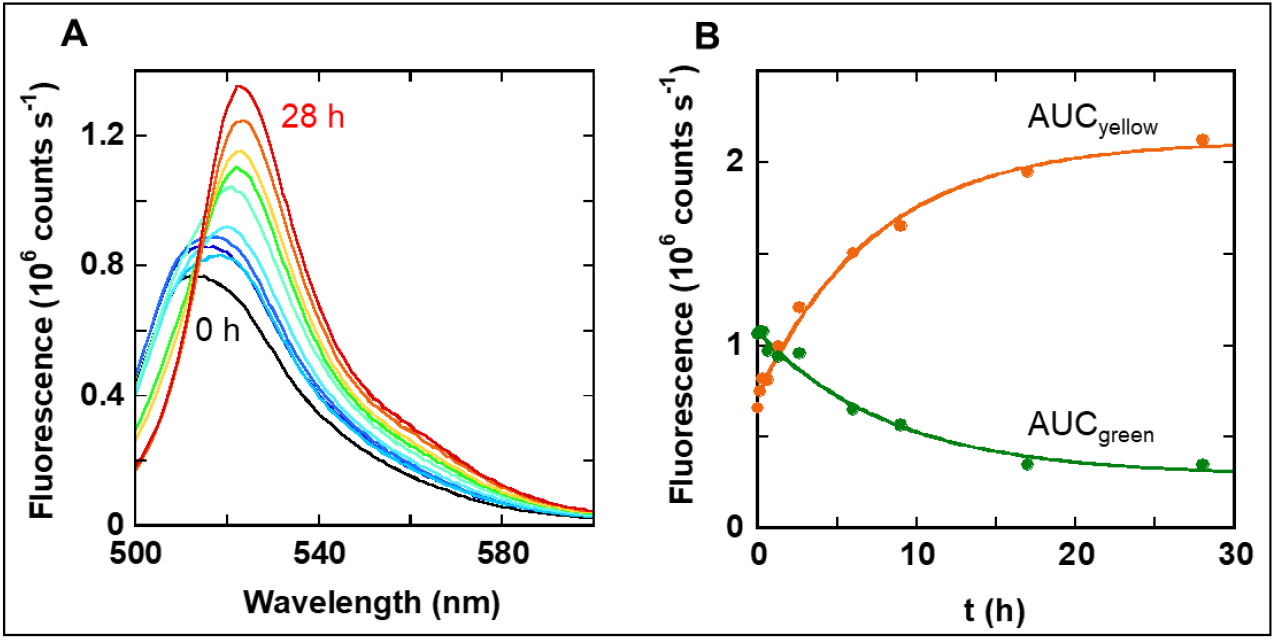
Sensor 2 is a ratiometric, ligand-induced color switch *in vitro*. (A) Unprocessed fluorescence spectra show a time-dependent shift from green to yellow emission after addition of 20 µM FK506 (45 °C). Spectra were collected at time intervals between t = 0 (black) and t = 28 h (red). (B) Ratiometric response is quantified by unmixing the spectra in panel A into AUC_yellow_ (orange) and AUC_green_ (green) components. Fitting these data to a single exponential function (lines) reveal t_1/2_ ∼ 5 h.

**Figure 3.**
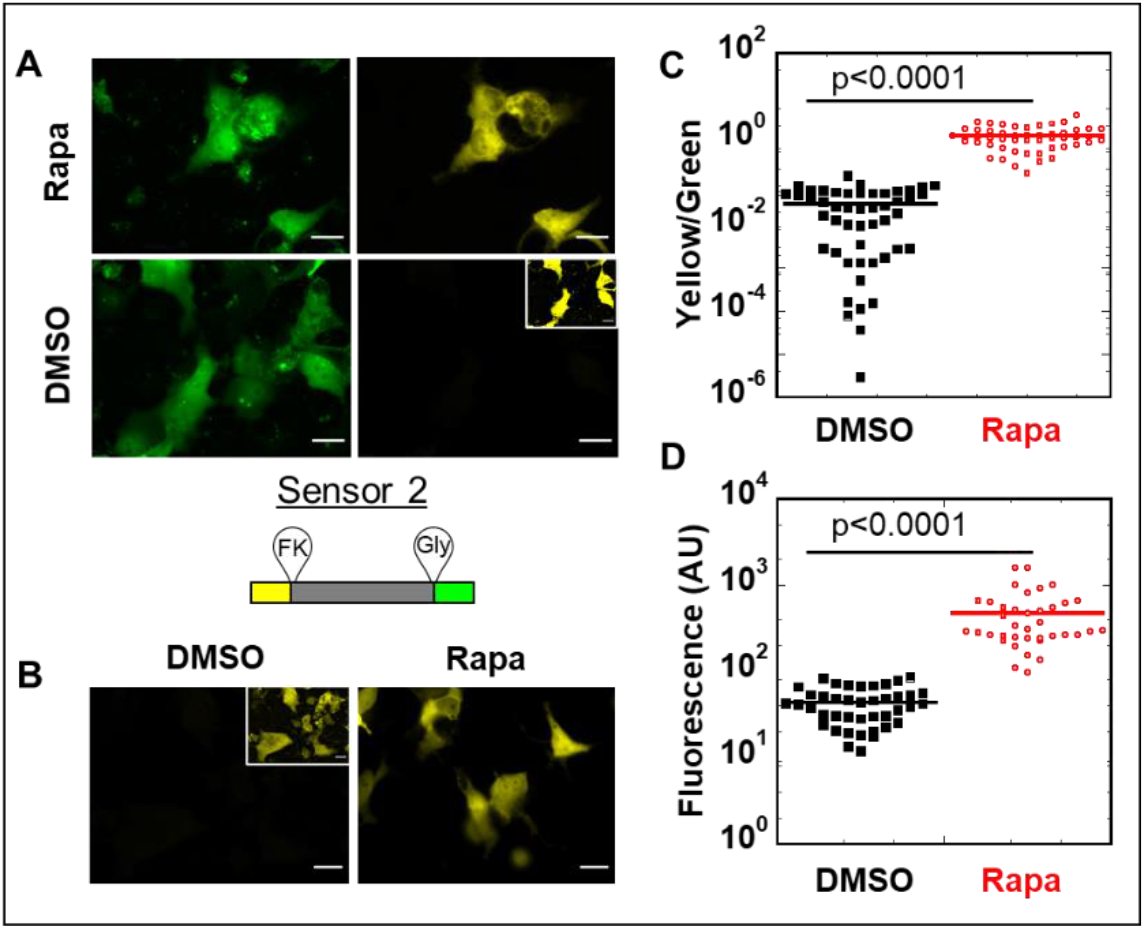
Sensor 2 and Y-fold analog show a large, intensiometric response in mammalian cells. (A) Representative fluorescence images of COS-7 cells expressing sensor 2 and treated with 5 µM rapamycin (top) or DMSO vehicle (bottom) for 4 h. Images were acquired in green and yellow channels and the relative contribution of G-fold and Y-fold was determined by unmixing the channels as described in Methods. Inset: the same image at higher contrast shows yellow fluorescence in cells treated with vehicle only, revealing that sensor 2 was expressed but only weakly fluorescent. (B) Representative images of COS-7 cells expressing Y-fold analog and treated with DMSO vehicle alone (left) or rapamycin (right). Inset: the same image in higher contrast shows that the Y-fold analog is expressed in cells but is only weakly fluorescent without rapamycin. (C) Sensor 2 turn-on was quantified by dividing the integrated cell intensity in yellow channel by the integrated cell intensity in the green channel. Each point represents a cell. Data are representative of three biological repeats as described in Methods (DMSO, *n* = 49; rapamycin, *n* = 51). Data for individual components of sensor 2 are shown in Figure S8. (D) Y-fold analog turn-on was quantified by integrating yellow fluorescence of cells expressing mKate2. Each dot represents a cell (DMSO, *n* = 38; rapamycin, *n* = 41). For panels C and D, significance was determined by performing a 2-tail Student’s t-test with unequal variance. Scalebars are 10 µm.

### Test of the entropy switch mechanism

The loop-closure entropy model holds that the Y-fold will be increasingly destabilized as the cpFKBP domain becomes more unstructured and flexible. To test this prediction, we unfolded and refolded sensor 2 as described above (45 °C), with inclusion of GdmCl in the refolding buffer to unfold any structure that remained in the cpFKBP domain. We chose concentrations of GdmCl (0.15 M and 0.3 M) that were sufficient to unfold free cpFKBP (Figure S2A) but too low to denature the cpFKBP-FK506 complex (Fig. S2B), the Y-fold analog, or the G-fold analog (Figure S1B). (Y/G)_OFF_ values decreased slightly from 0 M to 0.15 M GdmCl. This trend indicates that 0.15 M GdmCl destabilized the Y-fold relative to the G-fold, consistent with GdmCl unfolding residual structure in the cpFKBP domain. No further change in (Y/G)_OFF_ was observed with 0.3 M GdmCl, suggesting that cpFKBP was fully unfolded by 0.15 M GdmCl. Sensor 2 turn-on ratios and rates in 0.15 M and 0.3 M GdmCl were similar to the values without denaturant (Table 1).

In further support of the loop-closure entropy model, (Y/G)_OFF_ of sensor 2 progressively increased with decreasing temperature of refolding (45 °C, 37 °C, 22 °C; Figure S5D-F). This trend is likely caused by the cpFKBP domain folding at lower temperatures and thus stabilizing the Y-fold. Together, these data provide evidence for the esAFF mechanism *in vitro*.

### Sensor performance in cells

To test if our genetically-encoded biosensor functioned in mammalian cells, we transfected sensor 2 into COS-7 cells and treated the cells with rapamycin or DMSO vehicle. Sensor 2 was co-transfected with the red fluorescent protein mKate2, and cells exhibiting red fluorescence were randomly selected for imaging in green and yellow channels. We observed an 18-fold turn-on over DMSO vehicle alone within 30 min of adding rapamycin, with an average turn-on of (35 ± 9)-fold within four hours over three biological repeats (Figure 3A). The turn-on half-time was (60 ± 10 min) (Figure S8A). Sensor 2 responded to rapamycin in the expected dose-dependent manner (Figure S8B). Thus, sensor 2 performed exceedingly well in cells, achieving both a faster and a higher turn-on than what was attained *in vitro*.

Two observations, however, indicate that the turn-on mechanisms were different in cells and *in vitro*. Turn-on *in vitro* could largely be accounted for by the conversion of G-fold to Y-fold via AFF-mediated conformational switching. By contrast, green fluorescence in cells did not change significantly on rapamycin treatment (1.24 ± 0.53-fold). The change in (Y/G) was due almost exclusively to a large increase in yellow fluorescence (42 ± 23-fold), i.e., a dark-to-yellow process (Figure 3A, Figure S9). The second observation is that turn-on half-times were faster in cells (60 min) than *in vitro* (>4 h). These two findings suggest that turn-on arises from chromophore formation in the Y-fold in cells, and from the slower AFF-mediated conformational change *in vitro* (see Discussion).

We propose that the different sensing mechanisms *in vitro* and in cells originate from the putative kinetic trap that exists when the sensors fold in cells, together with the difference in protein expression temperatures, as follows. Protein samples for *in vitro* experiments were expressed at 18 °C in *E. coli*. At this temperature, sensor 2 expresses as a bright yellow protein because it is trapped in the Y-fold and folding as well as chromophore maturation are relatively efficient. Unfolding and refolding *in vitro* bypasses the kinetic trap, populating the G-fold and allowing the green-to-yellow fold shift to be observed. Sensor 2 was expressed in COS-7 cells at 37 °C. At this temperature, sensor 2 is mostly dark and becomes bright yellow in the presence of rapamycin. This result suggests that sensor 2 expresses mainly in the Y-fold, as it did at 18 °C, but the chromophore does not fluoresce until ligand is added. The half-time of 60 min suggests that turn-on is due to the slow process of chromophore maturation, not the comparatively rapid deprotonation of the already mature chromophore. If this view is correct, a likely reason why sensor 2 matures at 18 °C and not at 37 °C is that the cpFKBP domain is more disordered at the higher temperature. Lower temperatures (as well as FK506/rapamycin binding) stabilize cpFKBP, possibly tightening the Y-fold structure and facilitating chromophore maturation.

To test the hypothesis that sensor 2 turn-on in cells is due to the above mechanism and not the AFF-mediated fold shift, we transfected the Y-fold analog into COS-7 cells and asked whether removing the second copy of β10 changed the magnitude or rate of the yellow fluorescence increase. The Y-fold analog was also co-transfected with mKate2 and red fluorescent cells were randomly selected for imaging. Since the Y-fold analog has no green fluorescence, we calculated turn-on by the ratio of yellow intensities of rapamycin-treated vs. vehicle-treated cells. The Y-fold analog exhibited turn-on ((17 ± 7)-fold; Figure 3B) and t_1/2_ (∼50 min; Figure S8A) comparable to sensor 2, consistent with the chromophore maturation model. As a further test we attempted to recapitulate the COS-7 results in *E. coli* by expressing sensor 2 and the Y-fold analog at 37 °C. As expected, cell lysates exhibited only very weak yellow fluorescence (Figure S10). Addition of FK506 caused the yellow fluorescence to increase over several hours in the cell lysates, although not to the extent that was observed in COS-7 cells. No such increase was apparent when FK506 was added to protein expressed at 18 °C (Figure S1B). These findings argue that turn-on of sensor 2 (and the Y-fold analog) at 37 °C is primarily driven by ligand-assisted cpFKBP folding and subsequent chromophore maturation in the Y-fold.

## Discussion

We have demonstrated that inserting a binding domain into an engineered FP scaffold results in a ligand-driven, color-changing biosensor that operates according to the AFF mechanism *in vitro*. Several principles emerge as being important to the design of this class of switches. First, the thermodynamic stabilities of the two folds must be made comparable. In the current design, the β9-β10 and β10-β11 loops in GFP are amenable to inserting a ligand-binding domain and a disordered peptide for tuning the relative stabilities of the Y-fold and G-fold, respectively, without severe loss of fluorescence. The second consideration is to employ a receptor domain with an N-to-C distance that is compatible with the ∼9 Å distance between the C-terminus of β10 (S208) and the N-terminus of β11 (H217). There is likely some latitude in this distance requirement because we inserted the receptor between residues E213-K214 in the 8-amino acid loop that connects β10 and β11, and the residues to either side may act as somewhat flexible linkers. Many proteins naturally possess short (≤ 5 Å) N to C lengths^22^, and for those that don’t, circular permutation brings their new termini into proximity. Permutation sites can be identified using rational ^23^ or computational approaches^24^. Permutation typically destabilizes a protein, and if necessary further destabilization can be achieved by permuting using short linkers to pinch the original termini together ^25^, by introducing known mutations, or by chemical denaturants (Figure S6) and/or elevated temperature (Table S2).

Our study highlights how sensor behavior can change when it is expressed in cells versus refolded *in vitro*. The turn-on mechanism pivoted from an AFF-mediated green-to-yellow conformational change when sensor 2 was refolded, to dark-to-yellow chromophore maturation when it was expressed in mammalian cells. Although the structural features of the chromophore microenvironment and molecular details remain unclear, this finding emphasizes the roles that protein translation and temperature may play in folding and maturation of FPs. Of note, Boxer *et al*. reported that unfolding and refolding their related sensor also changed the relative populations of the two folds^19^.

Our AFF design blends elements of two main FP-based sensing mechanisms—bimolecular fluorescence complementation (BiFC) and that of genetically encoded calcium indicators (GECIs)—and offers the potential of increased modularity. In BiFC sensors, the FP is bisected into two complementary, nonfluorescent fragments that are attached to proteins that dimerize in response to ligand. An example is an N-terminal FP fragment fused to FKBP and the C-terminal FP fragment fused to FKBP-rapamycin binding protein (FRB)^26^. FKBP and FRB form a sandwich-type interaction with rapamycin. Dimerization increases the local concentration of the FP fragments, resulting in complementation and a turn-on that shares the same characteristics as sensor 2 and the Y-fold analog (i.e., dark-to-fluorescent and typically irreversible). BiFC sensors are straightforward in their construction but are not especially modular because the recognition domain consists of two heterologous proteins that must associate only in the presence of ligand. Finding natural protein partners that respond to a ligand of choice, or engineering proteins to do so, is often challenging. Additionally, the natural affinity that FP fragments have for one another makes turn-on dependent on sensor concentration, with spontaneous association and background fluorescence potential concerns.

GECIs are exemplified by the GCaMP family of calcium sensors^27^. In GCaMPs, the calmodulin (CaM) receptor domain and the M13/RS20 peptide are fused to an FP, at a point that abuts the chromophore^28^. Binding of Ca^2+^ to CaM causes it to associate with M13. This conformational change is transmitted to the FP, altering the local chemical environment around the chromophore and increasing its brightness. Sensor 2 behaves differently from GCaMPs in three respects. First, GCaMP chromophores mature in cells in the absence of ligand. Turn-on is consequently much faster than that of sensor 2 or the Y-fold analog, and it is readily reversible. The second distinction is that GCaMP sensors are typically monochromatic and their output is consequently intensity only. The readout of sensor 2 is intensiometric in cells but is ratiometric when the sensor is unfolded and refolded *in vitro*. The last difference is that GCaMPs, like BiFC sensors, require a specialized receptor domain that binds to a partner in the presence of the target analyte. The AFF design employs two principles—binding-induced folding and loop-closure entropy—that are more readily implemented into existing binding domains.

The main limitations with sensor 2 are its slow activation rate and that ratiometric output requires prior unfolding and refolding. Nevertheless, its 35-fold intensiometric turn-on in cells and 6-fold ratiometric response *in vitro* match those of other single FP-based genetically-encoded fluorescent sensors for Ca^2+ 29^, glucose^7^ and citrate^30^, and the GECO^31^ and GCaMP designs^32^. Due to the large fold-change and irreversible nature of their fluorescence turn-on, we envision that esAFF sensors (and their Y-fold analogs) may find utility in cellular applications in which one desires to detect targets that are present at very low concentration, where signal accumulation is advantageous.

A question that remains to be addressed is whether this design strategy can be applied to detect a range of useful biological targets in cells. In addition to its application in cellular sensing, the findings from this study can expand the protein engineers’ toolbox as it provides insight into how to create custom input and output functions by using a combination of protein engineering techniques: alternate frame folding of the output domain and circular permutation of the input domain. These characteristics confer modularity and may aid in the development of broadly applicable engineered protein switches.

## Methods

### Gene construction and protein purification

Amino acid sequences for all protein constructs are shown in Figure S10. Genes were cloned in a pET41b plasmid vector and fully sequenced. Proteins were expressed in *E*.*coli* BL21 (DE3) with isopropyl β-D-thiogalactopyranoside induction occurring at 18 °C for 16-18 h. Cell pellets were resuspended in 20 mM Tris (pH 8.0), 300 mM NaCl, 10 mM imidazole, 10 mM β-mercaptoethanol and lysed using a small amount of hen lysozyme followed by sonication. The soluble fraction was loaded onto a nickel-nitrilotriacetic acid column (Bio-Rad, Hercules, CA) and proteins were purified following the manufacturer’s protocols. Eluted proteins were dialyzed into buffer containing 10 mM Tris (pH 7.5), 150 mM NaCl. Each sensor construct was eluted as a monomeric species from a Superdex S200 Increase size exclusion column (Cytiva) and was judged to be > 95% pure by SDS-PAGE (Figure S5). The circularly permuted FKBP gene was inserted into the *Thermoanaerobacter tengcongensis* ribose binding protein gene as described by Ha *et al*.^33^ The cpFKBP-RBP fusion protein was expressed and purified as described above, after which RBP was cleaved off using HRV-3C protease. Free cpFKBP was recovered by passing the digested solution through a nickel-nitriloacetic acid column to remove free RBP and undigested cpFKBP-RBP.

### Spectroscopic measurements

Fluorescence spectra were recorded on a Horiba Jobin Yvon Fluoromax-4 benchtop fluorometer or a Molecular Devices Spectramax i3x plate reader. Settings for the benchtop fluorometer were 492 nm excitation (2 nm bandpass) and 500 – 600 nm emission (3 nm bandpass). Settings for the plate reader were 470 nm excitation and 500-600 nm emission. Refolding kinetics of cpFKBP was monitored by excitation at 280 nm (2 nm bandpass) and emission at 355 nm (6 nm bandpass).

### Protein stability determination

Solutions for denaturation experiments were prepared by mixing a solution of 20 mM Tris pH 8.0, 150 mM NaCl, 1 mM TCEP, 0.1 – 0.5 μM FP (or 3 μM cpFKBP) with an identical solution containing 1.0 M −4.4 M GdmCl, using a Hamilton Micro-lab 540B dispenser. Final denaturant concentrations were measured by refractive index. Samples were incubated for 24 h at 37 °C (FPs) or 8 h at room temperature (cpFKBP). Green/yellow fluorescence spectra of FP samples were recorded on the plate reader, and cpFKBP Trp emission data were collected on the benchtop fluorometer. GdmCl denaturation curves were fit to the linear extrapolation equation^34^ to obtain ΔG and m-values.

### Spectral unmixing

Fluorescence spectra were unmixed by fitting the data to Eq. 1 using MATLAB’s built-in nonlinear regression function (nlin-fit):

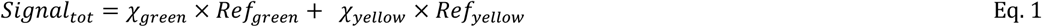

where *Signal*_*tot*_ is a 100-element vector consisting of the raw signal at each wavelength (500-600 nm), *Ref*_*green*_ and *Ref*_*yellow*_ are signals from the reference G-fold and Y-fold spectra, and *χ*_*green*_ and *χ*_*yellow*_ are scalars that correspond to the relative contributions of the G-fold and Y-fold. AUC_green_ and AUC_yellow_ were calculated by integrating *χ*_*green*_×*Ref*_*green*_ and *χ*_*yellow*_×*Ref*_*yellow*_, respectively.

Fluorescence microscopy images were unmixed in a similar way, except that *Signal*_*tot*_, *Ref*_*green*_, and *Ref*_*yellow*_ were vectors consisting of only two elements (the total signal from each pixel, the total signal of the G-fold, and the total signal of the Y-fold, in the green (excitation 470/40, emission 525/50) and yellow (excitation 500/20, emission 530/50) channels. We verified this method by mixing various ratios of purified G-fold and Y-fold analogs, imaging the solutions under the fluorescent microscope, and finding that the method accurately predicted their relative contributions.

### In vitro switching experiments

Sensors were unfolded at room temperature by adding 2/3^rd^ vol. of 50 mM HCl. After 2 min, the proteins were refolded by 1/30 dilution into experimental buffer (25 mM Tris (pH 7.5), 150 mM NaCl, 0.1 mM EDTA, 1 mM TCEP). The final protein concentration was ∼0.2 µM. The proteins were allowed to refold until all spectral changes were complete (1 h at 45 °C), at which point either 20 µM FK506 (ON state) or vehicle (0.04% DMSO; OFF state) were added. Fluorescence spectra were recorded on the Fluoromax-4 as described above. Turn-on values were calculated by dividing (Y/G)_ON_ by (Y/G)_OFF_ at the indicated time points.

To evaluate the possible contribution of a dark-to-yellow process to the observed turn-on, we first converted the unmixed fluorescence spectra to normalized concentrations of G-fold and Y-fold ([Gfold] and [Y-fold]) by dividing all AUC values by that of the G-fold in the OFF state and multiplying [Y-fold] values by 1.675 (the factor by which the Y-fold is brighter than the G-fold; Fig. 1C). After adding FK506, the decrease in [G-fold] (Δ[G-fold]) was calculated by [G-fold]_OFF_ – [G-fold]_ON_ and the increase in [Y-fold] (Δ[Y-fold]) was calculated by [Y-fold]_ON_ – [Y-fold]_OFF_. The percentage of turn-on due to the AFF mechanism is reported as Δ[Y-fold]/Δ[G-fold]. For *in vitro* experiments, these values were 100 % within experimental error, indicating that no dark-to-yellow component was present.

### COS-7 cell culture and imaging

Sensor 2 was transiently transfected under the constitutive CMV promoter and expressed for 36-48 h before adding rapamycin or DMSO vehicle in serum-free media. Turn-on was quantified by dividing the average intensity of each cell in the unmixed yellow channel by the average intensity in the unmixed green channel (Y/G). COS-7 cells were maintained in DMEM medium with 10% FBS, and Pen/Strep antibiotics in an incubator with 5% CO_2_ at 37 °C. Cells were split into fresh media one day before transfection in a six-well plate so that they reached 60-80% confluency the next day. Freshly prepared miniprep DNA (1.5 μg of pCMV-mKate2-N, and 1.5 μg of pCMV-MBP-sensor 2 or 1.5 μg pCMV-MBP-Y-fold analog) was then transfected using calcium phosphate method_35_, with the following modifications. CaCl_2_ was added to the plasmid mix to the final concentration of 250 mM (150 μL final volume), and 150 μL 50 mM HEPES 1.5 mM sodium phosphate (pH 7.0), 150 mM NaCl was then added dropwise while vortexing. The transfection mixture was added directly to media after aging at room temperature for 15 min, and the media was changed 12-15 h after transfection. 36 – 48 h after transfection, cells were washed with imaging media (DMEM, 25 mM HEPES pH 7.0, no FBS) and either rapamycin or DMSO vehicle was added to the cells diluted into the imaging media. Cells were treated with 5 μM rapamycin for 4 h to quantify the maximum turn-on, and for time points ranging from 30 min to 4 h to determine turn-on rate.

Cells were mounted in imaging media and imaged with Zeiss Axioimager Z1 upright microscope (40x/0.75 Plan-Neofluar objective) equipped with a mercury lamp, using the following filters: excitation 470/40, emission 525/50 for the green channel and excitation 500/20, emission 530/50 for the yellow channel. Cells were randomly picked using the mKate2 fluorescence. At least 10 fields were imaged for each replicate. For technical repeats (≥2 per sample), cells were split into a six-well dish on the same day and transfected from the same transfection mix, and for biological repeats (≥3 for each construct), cells were transfected by a fresh miniprep plasmid DNA on different days.

All cell images were processed using Fiji^36^. Background was subtracted from each channel using the built-in plugin, and the acquired image was then compressed. Each pixel of the resulting image was then unmixed to minimize the crosstalk between green and yellow channels using a custom MATLAB script as described above. Briefly, purified G-fold analog and Y-fold analog were imaged using the same settings to determine the crosstalk of each channel into the other, and this information was applied to linearly unmix the image of interest into their respective green and yellow contributions. Once unmixed, the intensity of each cell was measured in Fiji by manually drawing a region of interest around the periphery and determining the average intensity inside the region in both yellow and green channels. To quantify the turn-on for sensor 2, the yellow intensity for each cell was divided by the green intensity for that cell (Y/G), and for the Y-fold analog only the yellow channel was imaged and quantified.

## ASSOCIATED CONTENT

## AUTHOR INFORMATION

## ABBREVIATIONS

FP: fluorescent protein
cp: circular permutant
esAFF: entropy-switching alternate frame fold
GdmCl: guanidinium chloride
GECIs: genetically encoded calcium indicators
BiFC: bimolecular fluorescence complementation

