## Supporting Information for "Engineering A Fluorescent Protein Color Switch Using Entropy-driven Beta Strand Exchange"

**TABLE OF CONTENTS**

Figure S1. Spectral properties and stabilities of Y-fold and G-fold analogs.
Figure S2: Stability and refolding kinetics of cpFKBP.
Figure S3: Effect of unfolding/refolding and FK506 addition on sensor.
Figure S4. Destabilizing the G-fold by inserting residues at site B.
Figure S5. Purification and spectral characterization of sensor 2.
Figure S6: Effect of FK506 on sensor 2 under mildly denaturing conditions.
Figure S7. Sensor 2 shows lack of reversibility in the presence of WT FKBP.
Figure S8: Sensor 2 and Y-fold analog exhibits a rapamycin-induced intensiometric response in mammalian cells.
Figure S9. Sensor 2 responds to rapamycin predominantly via changes in yellow fluorescence, in mammalian cells.
Figure S10. Effect of FK506 on sensor 2 and Y-fold analog in cell lysates.
Figure S11. Amino acid sequence of proteins created for this study.

Table S1: Thermodynamic parameters for individual protein constructs at 37 ˚C
Table S2. Effect of temperature and GdmCl on the (Y/G) ratio of sensor 2 in the presence of FK506 (ON) or DMSO vehicle (OFF).

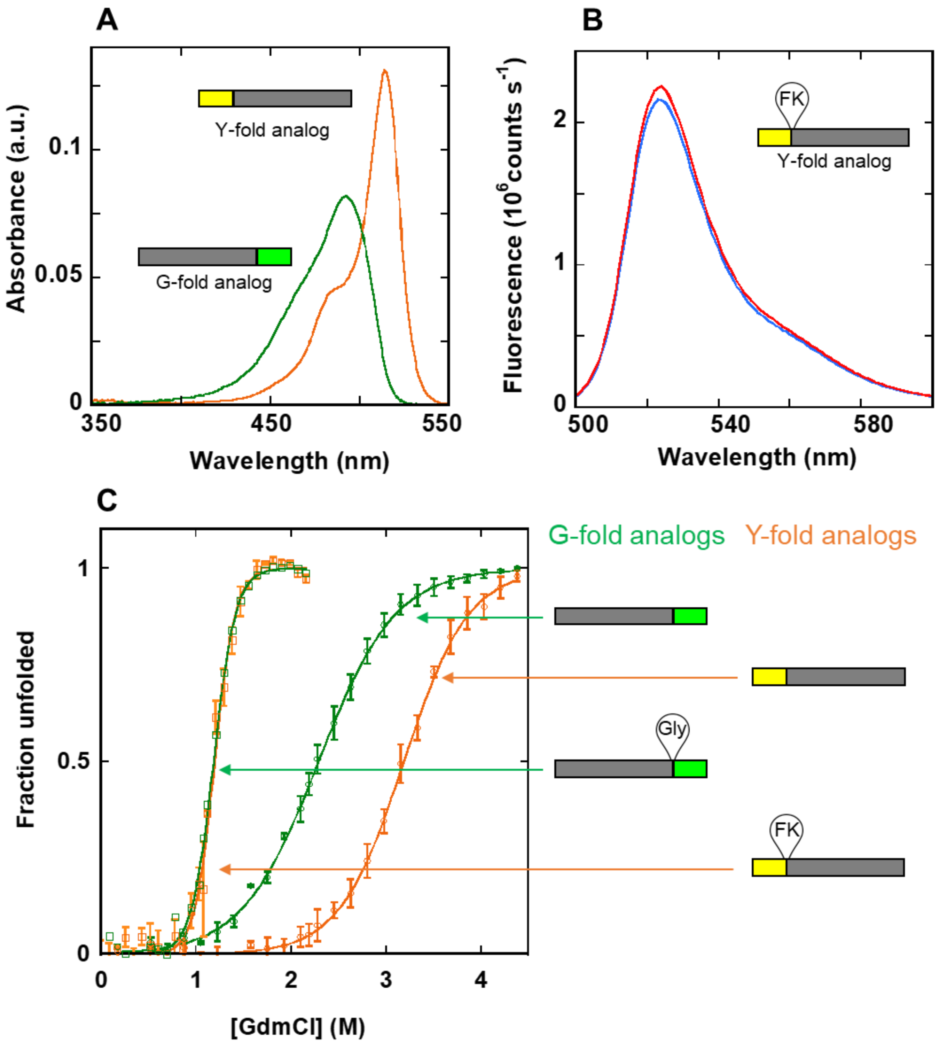


**Figure S1. Spectral properties and stabilities of Y-fold and G-fold analogs**. Insets in each panel show the position of strand β10 with 203Y (yellow) or a 203T (green). (A) Absorbance spectra of purified Y-fold and G-fold analogs are shown in orange and green, respectively. (B) Incubating purified Y-fold analog (expressed at 18 ˚C in *E. coli*) with FK506 (red) at 45 ˚C has no impact on its fluorescence relative to a DMSO control (blue) . (C) Guanidine denaturation of Y- and G-fold analogs of sensor 1 and sensor 2, obtained from fluorescence spectra and normalized to fraction unfolded ($n$ = 3, errors are SD). Orange and green filled circles represent Y-fold and G-fold, respectively, in the absence of insertions at site A or B (refer Figure 1B). Orange and green empty squares represent Y- and G-folds with a cpFKBP insertion and a single Gly insertion at site A and B, respectively.

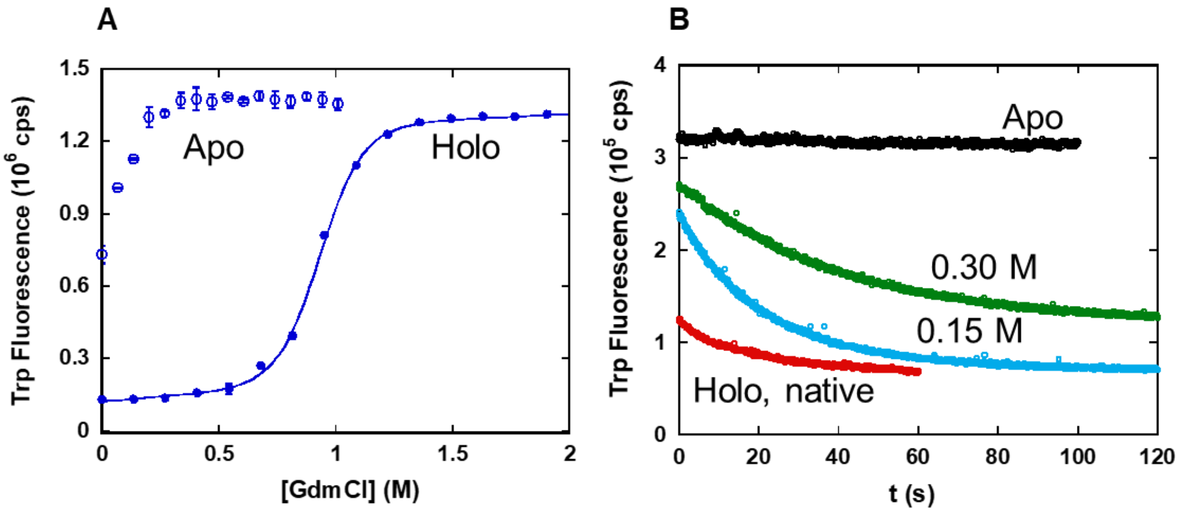


**Figure S2. Stability and refolding kinetics of cpFKBP.** (A) Guanidine denaturation curves of cpFKBP in the absence (open circles) and presence of FK506 (filled circles) recorded at 22 °C. Trp fluorescence was scanned after incubation of samples for 8 h. Fit parameters for the holo protein are: ΔG = 6.05 ± 0.15 kcal $mol^{-1}$, m = 6.49 ± 0.10 kcal $mol^{-1}$ $M^{-1}$, $C_m$ = 0.93 ± 0.01 M. Data represent average ± SD (*n* = 3). (B) FKBP induces refolding of cpFKBP as monitored by the decrease in Trp fluorescence. Unfolded cpFKBP (45 °C) is shown in black. Red, blue, and green curves were obtained by adding 20 μM FK506 to the unfolded protein in the presence of 0, 0.15 M, and 0.3 M GdmCl, respectively. Experimental buffer is 25 mM Tris pH 7.5, 150 mM NaCl, 0.1 mM EDTA.

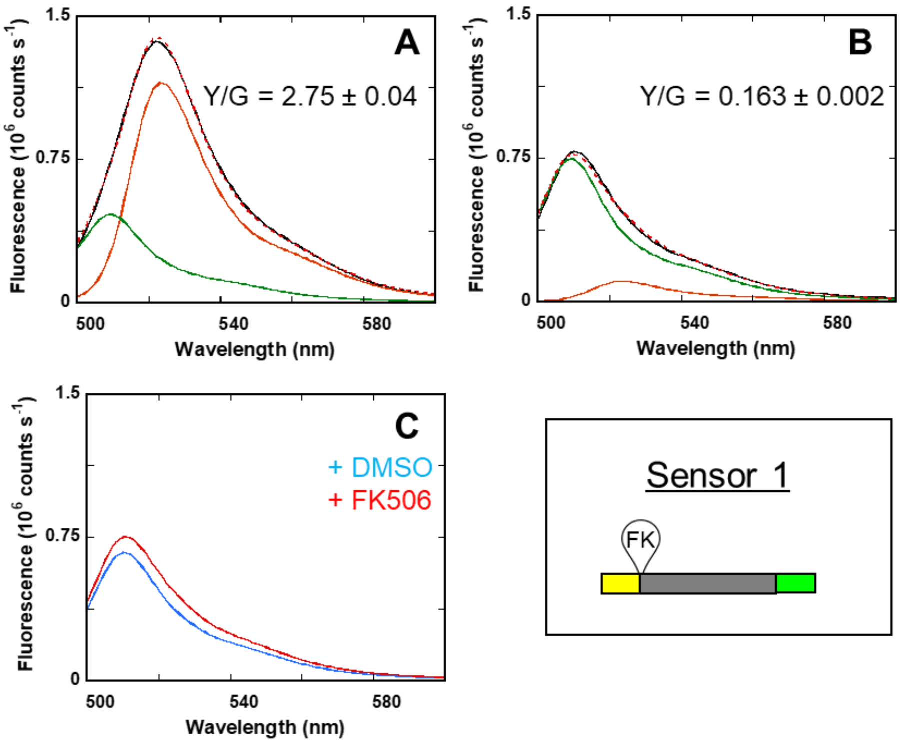


**Figure S3. Effect of unfolding/refolding and FK506 addition on sensor 1.** Fluorescence spectra of sensor 1 (A) before and (B) after unfolding/refolding indicate a large decrease in Y/G ratio after refolding *in vitro* at 45 °C. Raw spectra are shown in black, unmixed $AUC_{green}$ and $AUC_{yellow}$ curves in green and orange (respectively), and the sum of $AUC_{green}$ and $AUC_{yellow}$ in dashed red. (C) Addition of FK506/DMSO to refolded protein shown in (B) did not induce a spectral shift in sensor 1. Fluorescence spectra are shown with 20 μM FK506 (red) and 0.04 % DMSO vehicle (blue) after 4 hours of incubation with FK506 at 45 °C.

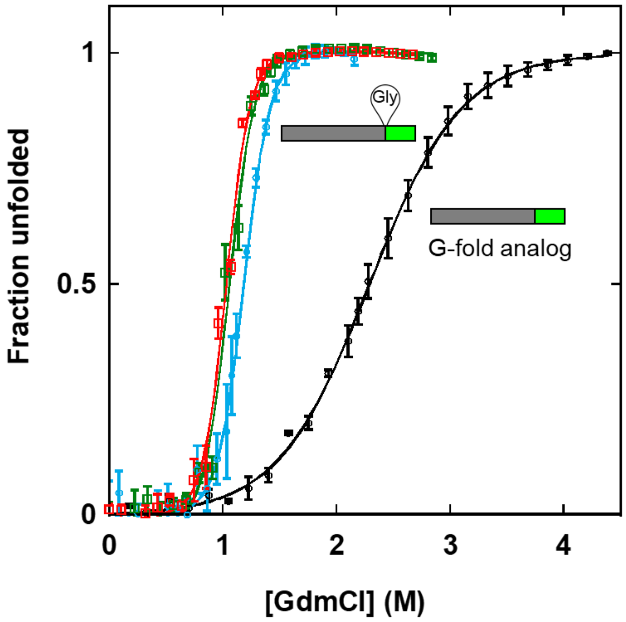


**Figure S4. Destabilizing the G-fold by inserting residues at site B.** $C_m$ (the [GdmCl] at which 50% protein is unfolded) of G-fold analog decreases from 2.27 ± 0.02 M to 1.18 ± 0.02 M with a single Gly insertion (blue) at site B (indicated in inset). Insertions of 5 (green) or 10 (red) residues at the same location didn't further decrease the $C_m$ significantly (n= 3, errors are SD). Fit parameters are reported in Table S1.

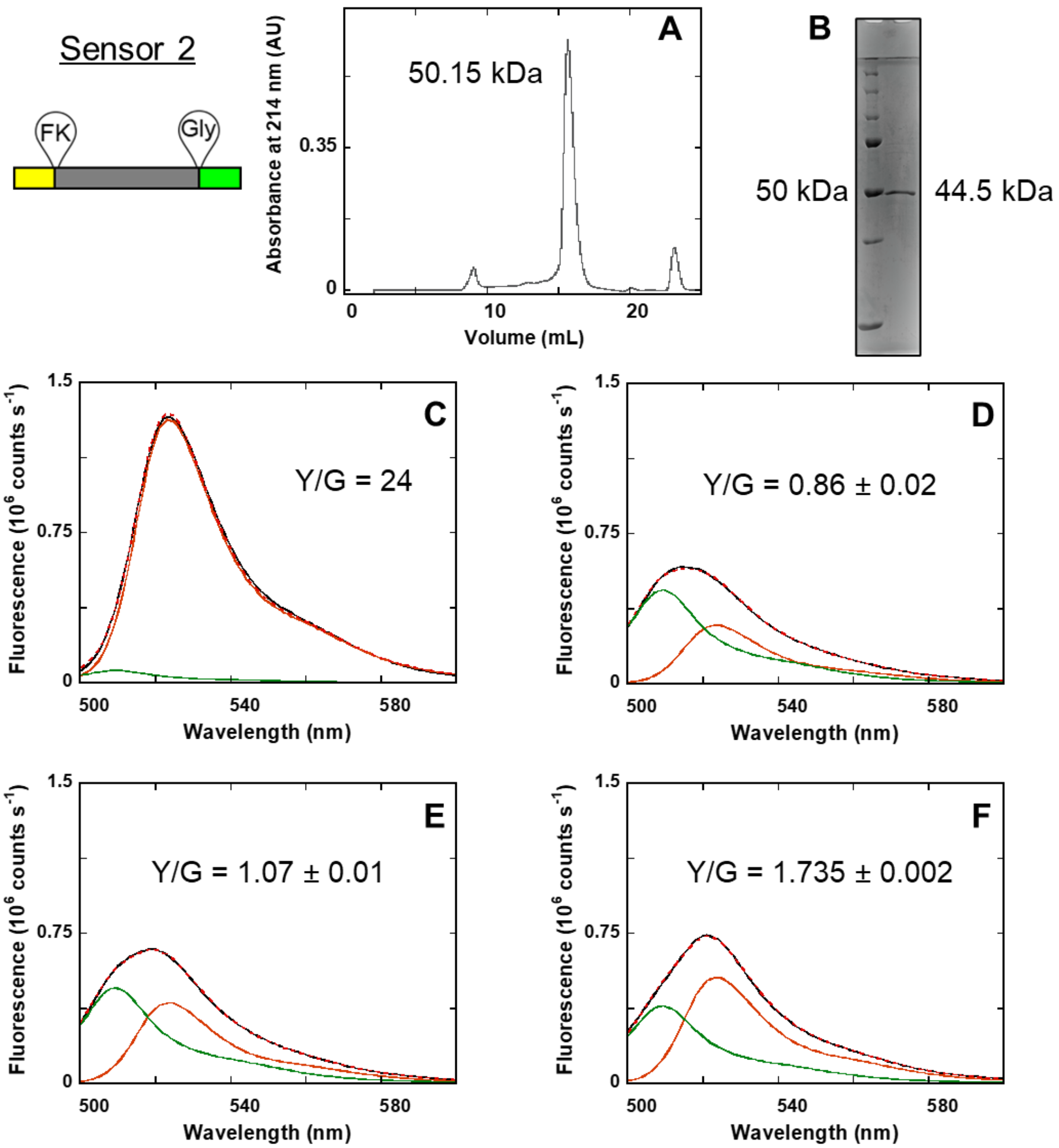


**Figure S5. Purification and spectral characterization of sensor 2**. (A) Size exclusion chromatogram of purified protein in 20 mM Tris pH 7.5, 150 mM NaCl, 0.005% Tween. (B) SDS-PAGE gel image with molecular weight marker on left and Sensor 2 purified on the nickel column shown on right. Sensor 2 (C) before, and after refolding *in vitro* under native conditions at (D) 45 ˚C (E) 37 ˚C and (F) 22 ˚C. The Y/G ratios are AUCs of the Y-fold and G-fold before incubating sensor 2 with DMSO/FK506 (as described in Methods). The data are average ± SD where *n* = 3 for D, E and F.

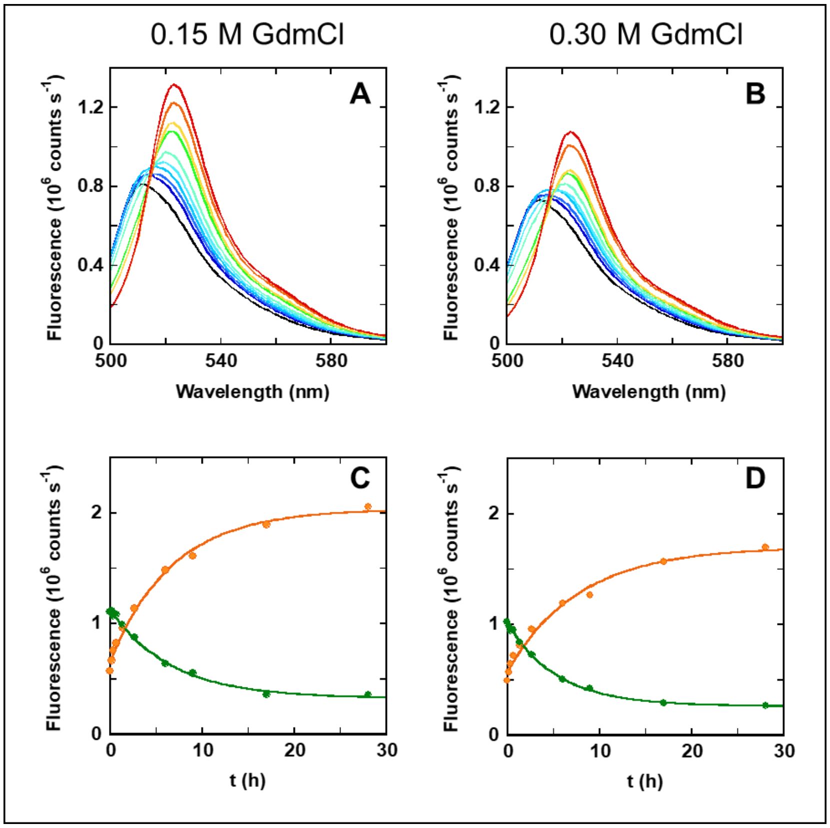


**Figure S6. Effect of FK506 on Sensor 2 under mildly denaturing conditions.** (A) and (B) represent spectral changes in response to FK506 binding cpFKBP in [GdmCl] < $C_m$ of the holo protein i.e., 0.93 M (refer Figure S2A). Spectra were collected at time intervals between t = 0 (black) and t = 28 h (red). (C) and (D) represent the time-dependent change in the unmixed areas under the curve (obtained as described in Methods), $AUC_{yellow}$ (orange) and $AUC_{green}$ (green) in 0.15 M and 0.3 M GdmCl, respectively.

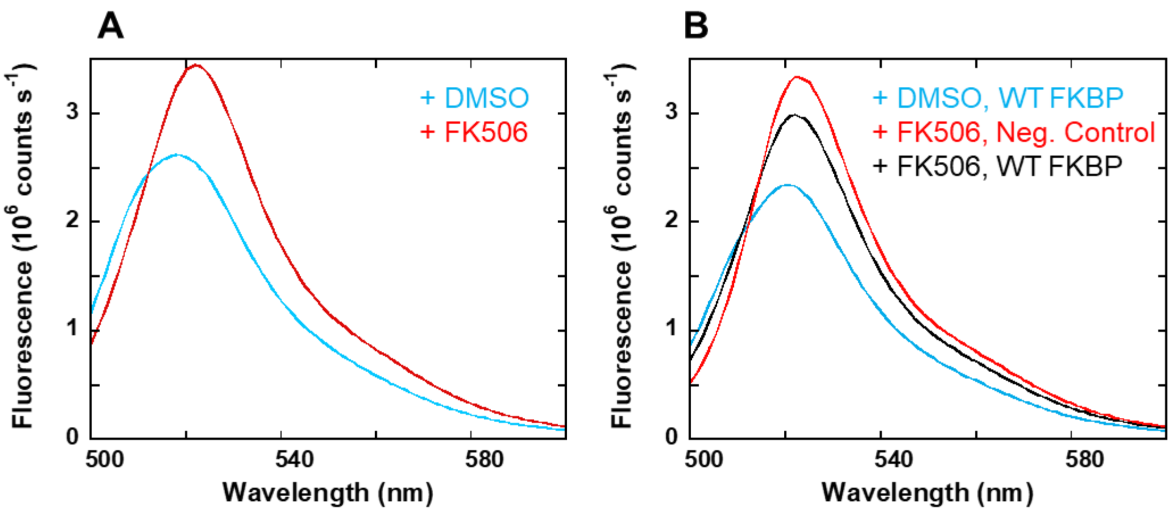


**Figure S7. Sensor 2 shows lack of reversibility in the presence of WT FKBP.** Spectra of (A) sensor 2 treated with (red) and without (apo, blue) 15 μM FK506 for 9 h at 45 ˚C. The respective Y/G ratios were 2.59 ± 0.24 and 0.954 ± 0.0005. (B) Each sample was treated with 30 μM WT FKBP or buffer (negative control). Black and red lines represent holo sensor 2 incubated with WT FKBP (Y/G = 2.51) and buffer (Y/G = 4.52) for 32 h. Blue line shows apo protein with WT FKBP (Y/G = 1.08). The Y/G ratios indicate that addition of WT FKBP prevented further change in Y/G but failed to reverse the reaction significantly in the time course of this experiment. Data are average ± SD, *n* = 2.

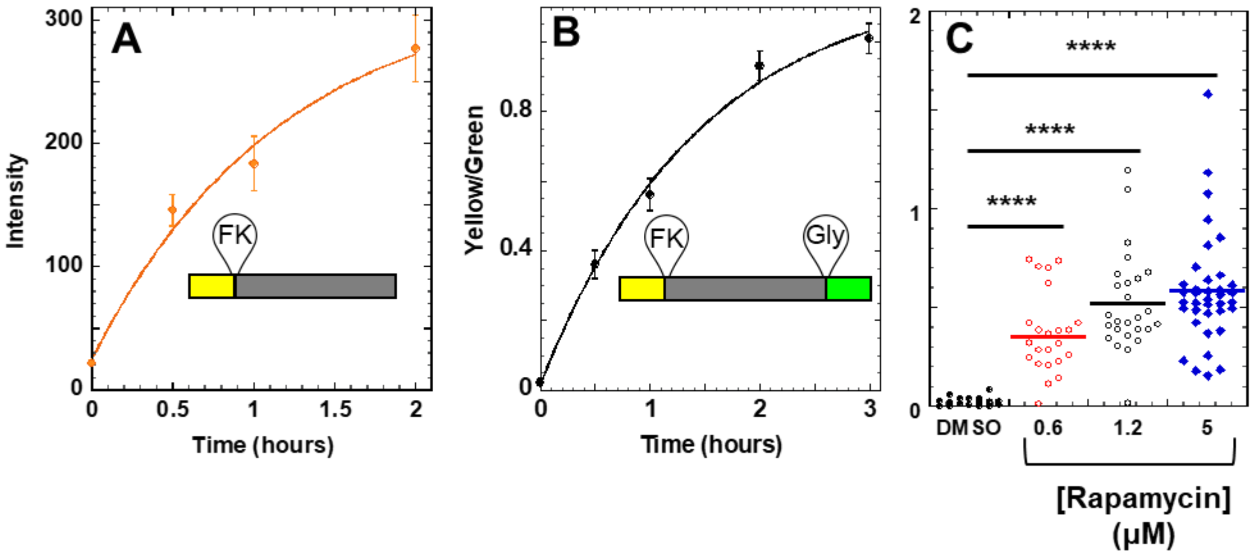


**Figure S8**. **Sensor 2 and Y-fold analog exhibits a rapamycin-induced intensiometric response in mammalian cells**. (A) Kinetics of Y-fold analog plotted as a function of change in fluorescence intensity observed in the yellow channel over 2 hours ($t_{1/2}$ = ~ 60 min). (B) Kinetics of sensor 2 plotted as a function of change in yellow:green (Y/G) ratios over 3 h ($t_{1/2}$ of 50 min ± 10 min. (C) Concentration dependent change in Y/G after 4 h of rapamycin exposure. Rapamycin concentrations at 0.6 μM (red), 1.2 μM (empty black circles) and 5 μM (filled blue circles) are shown with control, DMSO vehicle (black filled circles). **** = $p < .0001$ by Student t test for unpaired data with unequal variance and $n$ = 25, 23, 25, and 36 cells for DMSO, 0.6 μM ,1.2 μM, and 5 μM Rapamycin, respectively.

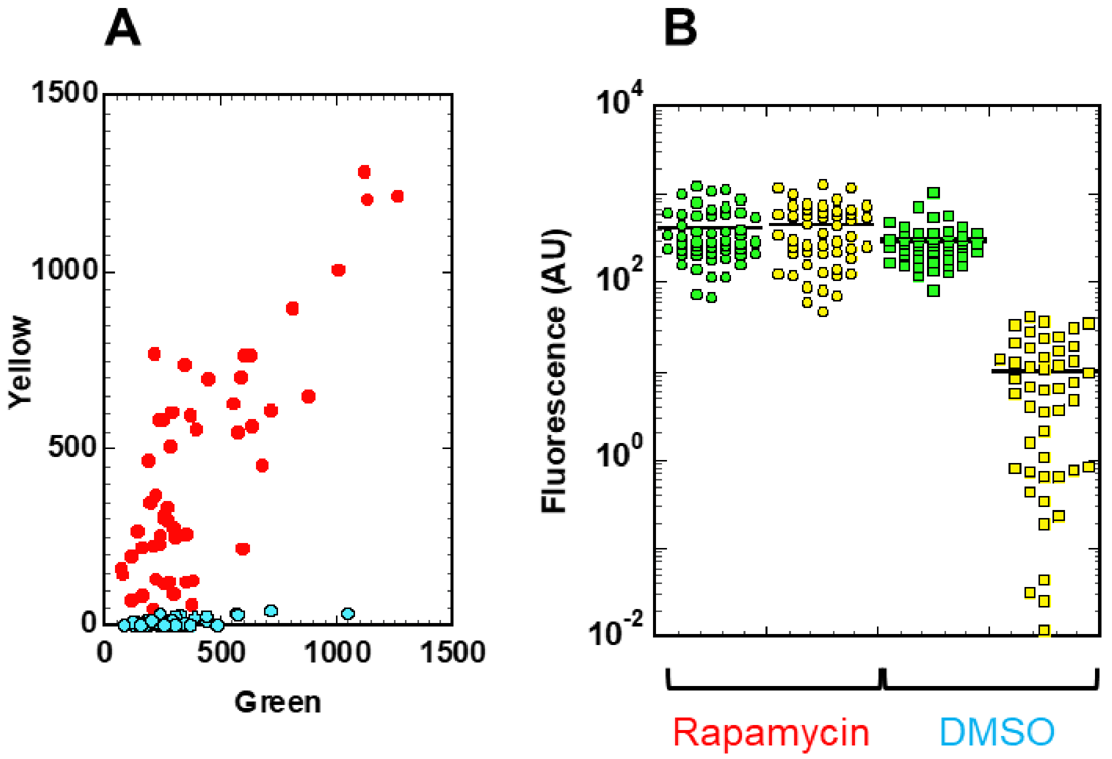


**Figure S9. Sensor 2 responds to rapamycin predominantly via changes in yellow fluorescence, in mammalian cells**. COS-7 cells were transfected with a plasmid encoding for sensor 2 and exposed to rapamycin or DMSO vehicle as described in methods. A) Raw green and yellow fluorescence in each analyzed cell (represented by dot) in presence of rapamycin (red) or DMSO alone (blue). B) Aggregated data showing green and yellow intensity of each cell either exposed to rapamycin or DMSO. Green and Yellow dots are representative of fluorescence in the green or yellow channel. Data is representative of 3 biological repeats. In all cases we saw a disproportionate change in yellow as compared to green.

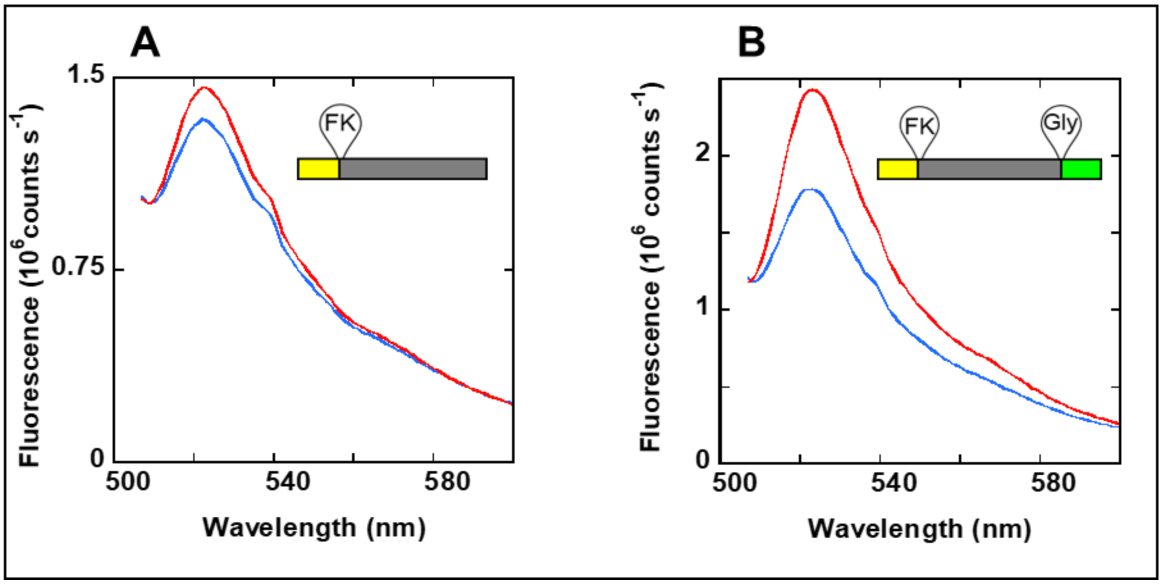


**Figure S10. Effect of FK506 on sensor 2 and Y-fold analog in cell lysates.** Post cell lysis emission spectra of (A) Y-fold and (B) sensor 2 expressed in *E.coli* at 37 ˚C, were collected after 8 h of treatment with 0.04% DMSO (blue) or 20 μM FK506 (red). Experimental buffer is 20 mM Tris pH 8.0, 300 mM NaCl and 100 mM EDTA. Insets represent protein constructs.

**Figure S11. Amino acid sequences of proteins created for this study.**

For sensor 1 and sensor 2, the N-terminal copy of strand β10 is orange, the N-terminal copy of strand β10 is green, the GYG chromophore is in italics, and the cpFKBP insertion is in blue. The color-specifying residue at position 203 (Tyr or Thr) is underlined. Linkers used in circular permutants are in bold.

1. G-fold analog of sensor 1
MKRDHMVLLEFVTAAGITLGMDELYK**GGGSGG**MVSKGEELFTGVVPILVELDGDVNGHKFSVRGEGEGDATNGKLTLKLICTT
GKLPVPWPTLVTTL*GYG*LACFSRYPDHMKQHDFFKSAMPEGYVQERTISFKDDGTYKTRAEVKFEGDTLVNRIELKGIDFKED
GNILGHKLEYNFNSHNVYITADKQKNGIKANFKIRHNVEDGSVQLADHYQQNTPIGDGPVLLPDNHYLSTQSVLSKDPNELEHH
HHHHHH

2. Circularly permuted FKBP
GKKFDSSRDRNKPFKFMLGKQEVIRGWEEGVAQMSVGQRAKLTISPDYAYGATGHPGIIPPHATLVFDVELLKLE**GGAASGGA
AGGSSGAASSGAGAAGGSGAGG**GVQVETISPGDGRTFPKRGQTAVVHYTGMLED

3. Y-fold analog of sensors 1 and 2
MLPDNHYLSYQSVLSKDPNEGKKFDSSRDRNKPFKFMLGKQEVIRGWEEGVAQMSVGQRAKLTISPDYAYGATGHPGIIPPH
ATLVFDVELLKLE**GGAASGGAAGGSSGAASSGAGAAGGSGAGGG**VQVETISPGDGRTFPKRGQTAVVHYTGMLEDKRDHM
VLLEFVTAAGITLGMDELYKGGGSGGMVSKGEELFTGVVPILVELDGDVNGHKFSVRGEGEGDATNGKLTLKLICTTGKLPVP
WPTLVTTL*GYG*LACFSRYPDHMKQHDFFKSAMPEGYVQERTISFKDDGTYKTRAEVKFEGDTLVNRIELKGIDFKEDGNILGHK
LEYNFNSHNVYITADKQKNGIKANFKIRHNVEDGSVQLADHYQQNTPIGDGPVLLEHHHHHHHH

4. Sensor 1
MLPDNHYLSYQSVLSKDPNEGKKFDSSRDRNKPFKFMLGKQEVIRGWEEGVAQMSVGQRAKLTISPDYAYGATGHPGIIPPH
ATLVFDVELLKLEGGAASGGAAGGSSGAASSGAGAAGGSGAGGGVQVETISPGDGRTFPKRGQTAVVHYTGMLEDKRDHM
VLLEFVTAAGITLGMDELYK**GGGSGG**MVSKGEELFTGVVPILVELDGDVNGHKFSVRGEGEGDATNGKLTLKLICTTGKLPVP
WPTLVTTL*GYG*LACFSRYPDHMKQHDFFKSAMPEGYVQERTISFKDDGTYKTRAEVKFEGDTLVNRIELKGIDFKEDGNILGHK
LEYNFNSHNVYITADKQKNGIKANFKIRHNVEDGSVQLADHYQQNTPIGDGPVLLPDNHYLSTQSVLSKDPNELEHHHHHHHH

5. G-fold analog of sensor 2

MKRDHMVLLEFVTAAGITLGMDELYK**GGGSGG**MVSKGEELFTGVVPILVELDGDVNGHKFSVRGEGEGDATNGKLTLKLICTT
GKLPVPWPTLVTTL*GYG*LACFSRYPDHMKQHDFFKSAMPEGYVQERTISFKDDGTYKTRAEVKFEGDTLVNRIELKGIDFKED
GNILGHKLEYNFNSHNVYITADKQKNGIKANFKIRHNVEDGSVQLADHYQQNTPIGDGPVLGLPDNHYLSTQSVLSKDPNELEH
HHHHHHH

6. Sensor 2
MLPDNHYLSYQSVLSKDPNEGKKFDSSRDRNKPFKFMLGKQEVIRGWEEGVAQMSVGQRAKLTISPDYAYGATGHPGIIPPH
ATLVFDVELLKLEGGAASGGAAGGSSGAASSGAGAAGGSGAGGGVQVETISPGDGRTFPKRGQTAVVHYTGMLEDKRDHM
VLLEFVTAAGITLGMDELYK**GGGSGG**MVSKGEELFTGVVPILVELDGDVNGHKFSVRGEGEGDATNGKLTLKLICTTGKLPVP
WPTLVTTLGYGLACFSRYPDHMKQHDFFKSAMPEGYVQERTISFKDDGTYKTRAEVKFEGDTLVNRIELKGIDFKEDGNILGHK
LEYNFNSHNVYITADKQKNGIKANFKIRHNVEDGSVQLADHYQQNTPIGDGPVLGLPDNHYLSTQSVLSKDPNELEHHHHHHHH
H

**Table S1. Thermodynamic parameters for individual protein constructs at 37 ˚C**

| **Fluorescent protein constructs** | **ΔG (kcal mol-1)** | **m (kcal mol-1 M-1)** | **Cm (M)** |
|---|---|---|---|
| Y-fold (no cpFKBP insertion) | 6.49 ± 0.62 | 2.11 ± 0.15 | 3.07 ± 0.08 |
| Y-fold analog (cpFKBP insertion) | 5.32 ± 0.32 | 4.29 ± 0.29 | 1.24 ± 0.01 |
| G-fold analog | 3.71 ± 0.05 | 1.64 ± 0.02 | 2.27 ± 0.02 |
| G-fold analog (Gly insertion) | 6.38 ± 1.49 | 5.4 ± 1.19 | 1.18 ± 0.02 |
| G-fold analog (GSGSG insertion) | 5.81 ± 0.12 | 5.53 ± 0.20 | 1.05 ± 0.02 |
| G-fold analog (GGSGTSGGSG insertion) | 6.15 ± 0.63 | 5.97 ± 0.53 | 1.03 ± 0.01 |

GdmCl-induced unfolding of fluorescent proteins was monitored by fluorescence emission maxima (524 nm and 509 nm for Y-fold and G-fold, respectively). Thermodynamic parameters correspond to the graphs shown in Figures S1 and S4. Data represent average ± SD ($n$ = 3).

**Table S2. Effect of temperature and GdmCl on the (Y/G) ratio of sensor 2 in the presence of FK506 (ON) or DMSO vehicle (OFF)**

| Conditions (Temperature, [GdmCl]) | $(Y/G)_{OFF}$ | $(Y/G)_{ON}$ |
|---|---|---|
| 45 ˚C, 0 M[a] | 1.15 ± 0.25 | 6.62 ± 0.69 |
| 45 ˚C, 0.15 M[a] | 0.94 ± 0.14 | 6.10 ± 1.08 |
| 45 ˚C, 0.3 M[b] | 0.93 ± 0.14 | 5.77 ± 0.35 |
| 37 ˚C, 0 M[a] | 1.77 ± 0.06 | 1.81 ± 0.01 |
| 22 ˚C, 0 M[a] | 3.77 ± 0.25 | 2.15 ± 0.02 |

$(Y/G)_{OFF}$ and $(Y/G)_{ON}$ denote the (Y/G) values of sensor 2 at the final time point recorded for each condition with 0.04 % DMSO and 20 μM FK506, respectively. (Y/G) values were calculated from AUCs of Y-fold and G-fold analogs (as described in Methods) at the respective temperatures and buffer conditions. [a]Data are average ± SD (*n* = 3). [b]Data are average ± SD (*n* = 2).
